# Preovulatory oocyte aging in mice affects fertilization rate and embryonic genome activation

**DOI:** 10.1101/209437

**Authors:** Hannah Demond, Debora Dankert, Ruth Grümmer, Bernhard Horsthemke

## Abstract

Delayed ovulation, or preovulatory aging, can seriously compromise the developmental competence of oocytes. In the present study, we have investigated the effect of preovulatory aging on preimplantation embryos. Delaying ovulation with the gonadotropin releasing hormone (GnRH) antagonist Cetrorelix led to a decline in 2-cell rate from 76 to 46%. From control mice, an average of 17 embryos per mouse was retrieved. This number decreased to a mean of 5 embryos per mouse after preovulatory aging, suggesting that fertilization is impaired by aging. For analysis of zygotic genome activation, 2-cell embryos were incubated with BrUTP, which was incorporated into nascent RNA and detected by immunohistochemistry. A 2.85-fold increase in fluorescence intensity was detected after aging, pointing to a precocious activation of the genome. A possible effect of preovulatory aging on genomic imprint maintenance was investigated at the 8-cell stage. Deep amplicon bisulfite sequencing of *Igf2r*, *Snrpn*, *H19* and *Pou5f1* showed no significant changes between embryos derived from preovulatory-aged oocytes and control embryos, indicating stable imprint maintenance throughout epigenetic reprogramming. We conclude that preovulatory aging of the oocyte affects fertilization and embryonic genomic activation.

## Introduction

Oocyte maturation, fertilization and early embryonic development are highly regulated processes and their synchronized timing has major implications for further development and health later in life. Delayed ovulation, also known as preovulatory aging, leads to impairment of embryonic progression in several animal species (Smits *et al.* 1995). In humans, a delay in ovulation may occur due to cycle irregularities (Smits *et al.* 1995) or in the course of assisted reproductive techniques when hormonal treatment is prolonged. It is associated with decreased implantation potential, malformations, chromosomal abnormalities, decreased embryonic weight and a high mortality rate (Smits *et al.* 1995; Bittner *et al.* 2011). The molecular causes behind the decreased oocyte competence of preovulatory-aged oocytes are still unclear. In a previous study we showed that preovulatory aging affects the expression levels and poly(A)-tail length of maternal effect genes (Dankert *et al.* 2014). This group of genes is expressed in the oocyte but regulates processes in the early embryo such as zygotic genome activation (ZGA) and the maintenance of DNA methylation of imprinted genes. The deregulation of maternal effect genes after preovulatory aging is likely to affect the developmental potential of the oocyte.

In early embryonic development, ZGA is required for the maternal-to-embryonic transition, which includes the replacement of maternal effect gene mRNAs and proteins from the oocyte by newly synthesized embryonic factors. In mice, transcription is activated in several consecutive waves. The first major step occurs around the 2-cell stage and is crucial for further embryonic development (Schultz 1993; Hamatani *et al.* 2004; Wang *et al.* 2004). In the preovulatory-aged oocyte, relative mRNA expression of *Smarca4*, *Tet3* and *Zfp57* is decreased (Dankert *et al.* 2014). *Smarca4* encodes the catalytic subunit of SWI/SNF-related complexes and regulates chromatin remodeling in the process of ZGA (Bultman *et al.* 2006). TET3 protein oxidizes 5-methylcytosine (5mC) to 5-hydroxymethylcytosine (5hmC) and is critical for global demethylation of the parental genomes during ZGA (Gu *et al.* 2011; Peat *et al.* 2014).

Some of the maternal effect genes encode proteins that are required for the maintenance of genomic imprints during epigenetic reprogramming in embryonic preimplantation development, e.g. *Zpf57* gene (Li *et al.* 2008; Messerschmidt 2012). Genomic imprinting is an epigenetic process by which the male and the female germline confer different DNA methylation marks onto specific gene regions, so that one allele of an imprinted gene is active and the other one is silent. It is of vital importance for normal embryonic and fetal development that the methylation imprints are protected against the waves of genome-wide DNA demethylation and re-methylation that occur before and after the blastocyst stage, respectively. In humans, loss of differential methylation at imprinted genes leads to severe imprinting syndromes, e.g. Beckwith-Wiedemann and Angelman syndrome (Horsthemke and Wagstaff 2008; Eggermann 2009).

To analyze whether the deregulation of maternal effect genes observed after preovulatory aging (Dankert *et al.* 2014) affects the developmental potential of the oocyte, the current study investigated the developmental competence of preovulatory-aged oocytes. Fertilization rates as well as the capacity to activate the embryonic genome at the 2-cell stage were found to be affected by preovulatory aging. Furthermore, we analyzed DNA methylation levels in single 8-cell embryos by deep amplicon bisulfite sequencing of 3 imprinted genes (*H19*, *Snprn*, *Igf2r*) and *Pou5f1*, an unmethylated control gene that encodes the OCT4 protein, to investigate a possible influence of preovulatory oocyte aging on imprinting in the early embryo. DNA methylation appeared stable in embryos from preovulatory-aged oocytes.

## Results

### Impaired fertilization after preovulatory aging

Preovulatory aging was induced in C57Bl/6J female mice by delaying ovulation for 3 days, followed by mating with C57Bl/6J × CBA/Ca F1 hybrid males. In control mice 10 of 18 matings were successful, whereas after preovulatory aging embryos were found only in 5 out of 16 matings. Two-cell embryos were retrieved from the oviduct 36 hours after mating. On average, control mice yielded 16.6 ± 3.6 2-cell embryos (*n* = 10; Fig 1A). After preovulatory aging the embryo number dropped to a mean of 5.2 ± 2.5 embryos per mouse compared to controls (*n* = 5; *t*-test: *p* = 0.06). The lower number of embryos after preovulatory aging might be due to a decrease in oocyte number, since we previously described a decline in ovulation rates after preovulatory aging (Demond *et al.* 2016). To investigate whether fertilization rate is also affected by preovulatory aging, the ratio between 2-cell embryos and unfertilized oocytes flushed from the oviduct was calculated. The mean 2-cell embryo rate of control mice was 76.0% ± 6.9 and decreased significantly after preovulatory aging to 46.1% ±10.5 (Fig 1B; *t*-test *p <* 0.05). Taken together, it seems that preovulatory aging lowers the number of ovulated oocytes and that these oocytes are more difficult to fertilize, resulting in a decreased embryo-yield.

**Fig 1:**
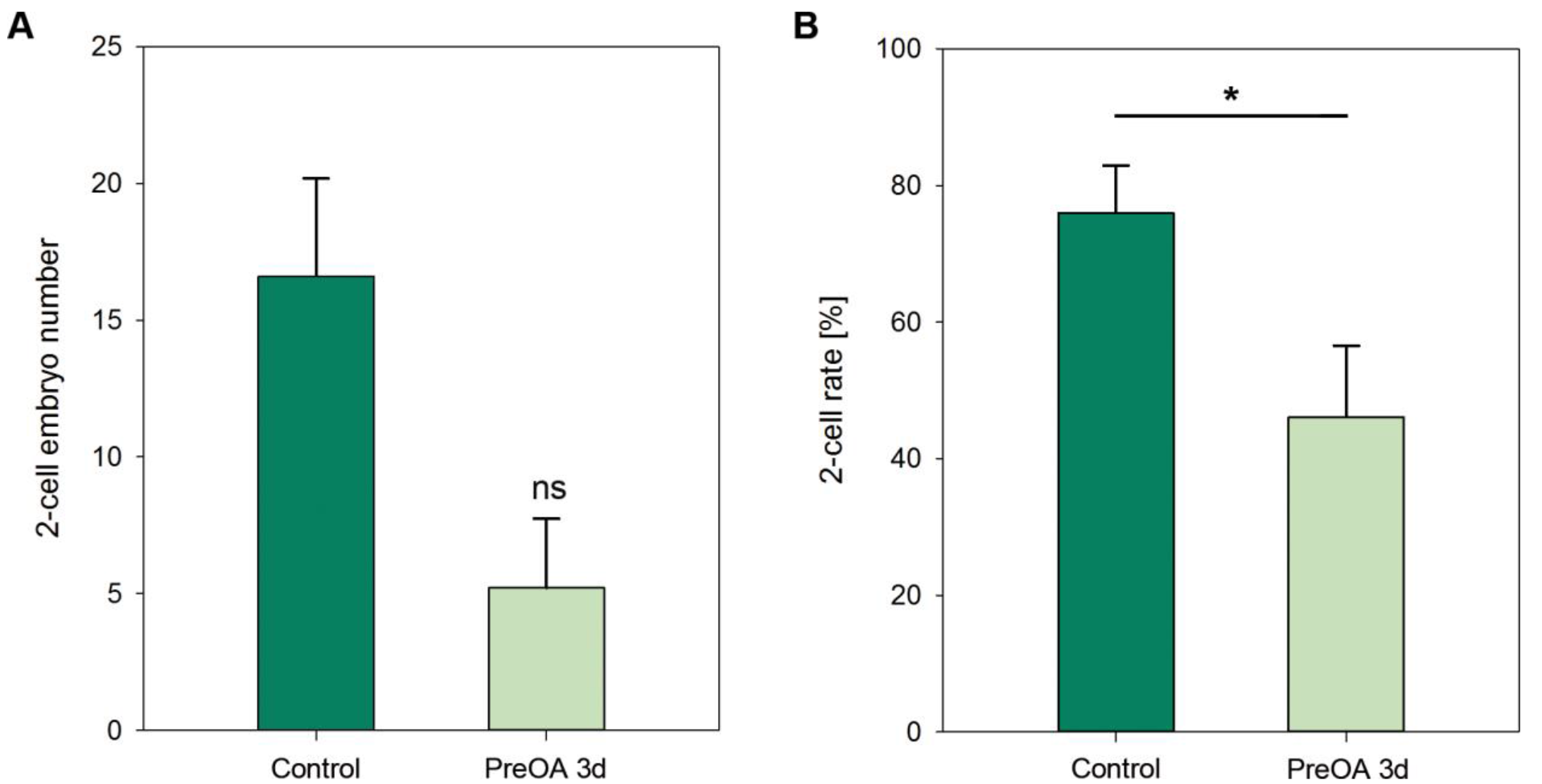
Impaired fertilization of preovulatory-aged (PreOA) oocytes compared to control oocytes. Control and preovulatory-aged oocytes from 5 and 9 mice respectively, were fertilized by natural mating after postponing ovulation for 3 days. **A)** After preovulatory aging less 2-cell embryos per mouse were retrieved and **B)** 2-cell embryo rate decreased significantly compared to controls (Mann-Whitney U: n.s. > 0.05; \**p*< 0.05).

### BrUTP incorporation in 2-cell embryos after preovulatory aging shows increased transcription during embryonic genome activation

The relative amount of BrUTP incorporation in 2-cell embryos derived from preovulatory-aged and control oocytes was compared as indicator for transcription during zygotic genome activation. A total of three BrUTP incorporation experiments was carried out, each with embryos from 4 mice per group, which resulted in a total number of 60 control 2-cell embryos and 38 embryos derived from preovulatory-aged mice.

BrUTP signal was detected in the cell nuclei and nucleoli of control and preovulatory-aged 2-cell embryos (Fig 2A). The BrUTP signal appeared stronger in embryos derived from preovulatory-aged oocytes than in controls. Quantification of the fluorescent signal confirmed a significant 2.85 ± 0.30-fold increase of BrUTP signal after preovulatory aging compared to controls (Mann-Whitney U: *p* < 0.001; Figu 2B). This demonstrates an increase in transcription after preovulatory aging at the 2-cell stage of embryonic development.

**Fig 2:**
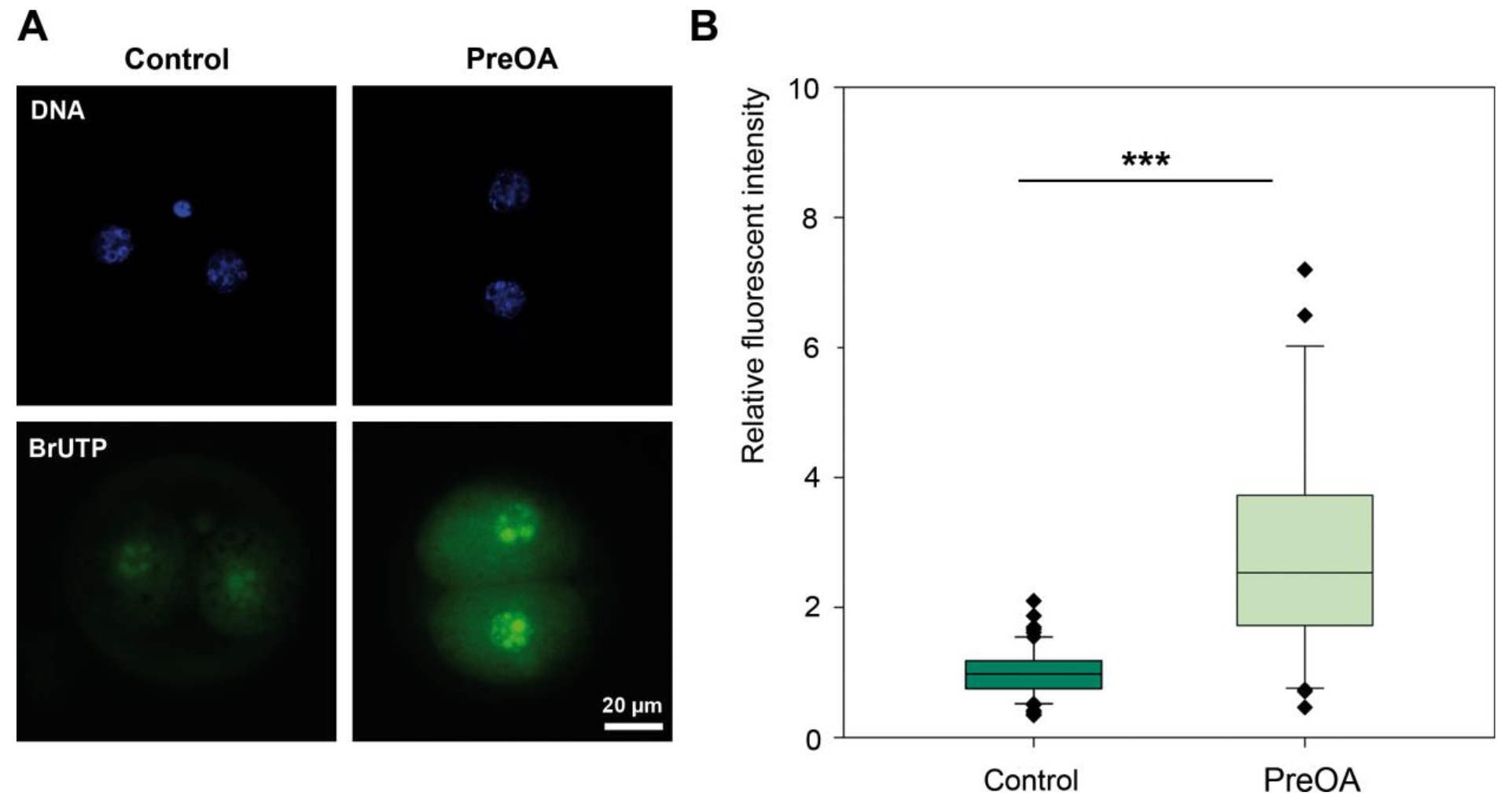
Immunofluorescence staining and quantification of incorporated BrUTP in 2-cell embryos derived from control and preovulatory-aged (PreOA) oocytes. **A)** DAPI incorporation (DNA) and anti-BrdU staining (BrUTP) of representative embryos, showing BrUTP incorporation in the cell nucleus. **B)** Quantification of fluorescence intensity demonstrates a significant increase in transcription after preovulatory aging compared to controls (Mann-Whitney U: \*\*\**p* < 0.001).

### Effects of preovulatory aging on DNA methylation levels in 8-cell embryos

DNA methylation levels of the three imprinted genes *Igf2r*, *Snrpn* and *H19* and the unmethylated control gene *Pou5f1* were determined in single 8-cell embryos by deep amplicon bisulfite sequencing. Embryos were obtained from 3 female mice for each group. In total, 15 control embryos and 14 embryos derived from preovulatory-aged oocytes were compared. Methylation levels for individual embryos for each CpG are listed separately for the maternal and paternal allele in supplementary figures S1-8. The *H19* amplicon included 4 CpGs but the methylation on the paternal allele of the first CpG was unstable in both control and preovulatory-aged embryos, so it was excluded from the analysis. The mean methylation levels of the paternal and maternal allele of each embryo were calculated. Methylation levels were approximately 50% for all analyzed imprinted genes and no significant differences were detected between two groups (Fig 3 A-C; *Igf2r*: Control = 47.2% ± 0.7, PreOA = 47.6% ± 0.5; *Snrpn*: Control = 42.5% ± 0.9, PreOA = 49.0% ± 0.8; *H19*: Control = 49.6 ± 0.6%, PreOA = 47.5% ± 0.6). The *Pou5f1* locus showed hypomethylation for both control embryos (5.0% ± 0.5) and embryos after preovulatory aging (1.6% ± 0.4; Fig. 3 D), revealing no statistical significant differences between the two groups. A post-hoc power analysis demonstrated a probability of 94.9% to detect a 10% methylation difference with the experimental set-up used. A difference of less than 10% was assumed not to be of biological relevance. The power analysis supports the results, demonstrating stable imprint maintenance in embryos up to the 8-cell stage after preovulatory aging for the loci analyzed in this study.

**Fig 3:**
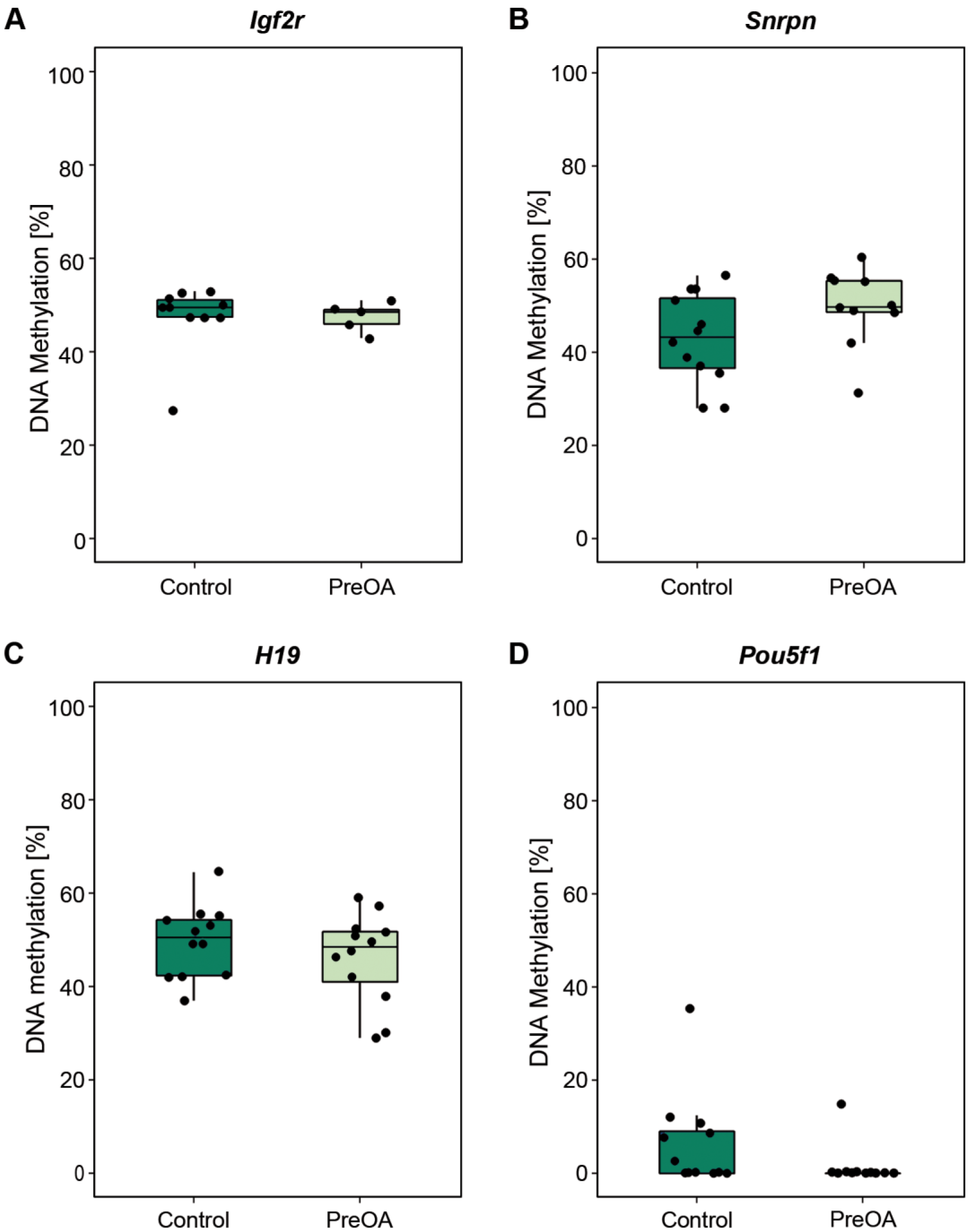
DNA methylation levels of 8-cell embryos derived from control and preovulatory-aged (PreOA) oocytes. Deep amplicon bisulfite sequencing of single embryos was used to analyze DNA methylation status of the maternally methylated imprinted genes, *Igf2r* **(A)** and *Snrpn* **(B)**; the paternally methylated imprinted gene *H19* **(C)** and the unmethylated *Pou5f1* locus **(D)**. No significant effect of preovulatory aging on DNA methylation levels was observed. Methylation status of single embryos is indicated by the black dots.

## Discussion

Preovulatory aging of murine oocytes is known to disturb postimplantation embryonic development (Bittner *et al.* 2011). In the current study we analyzed if also the development of the early preimplantation embryo is affected and found that preovulatory aging decreased 2-cell embryo rates significantly. The total number of embryos derived from aged oocytes was even lower than what would have been expected from the decreased fertilization rates. This is likely due to the decrease in ovulation rates after preovulatory aging (Demond *et al.* 2016). In rats, low fertilization rates after preovulatory aging have been associated with altered hormone levels, chromosomal abnormalities and altered RNA synthesis (Fugo and Butcher 1966; Peluso and Butcher 1974). Impaired RNA dynamics have also been described for murine oocytes (Dankert *et al.* 2014). Furthermore, chromosomal abnormalities were found after in vitro preovulatory aging in a follicle culture system (Demond *et al.* 2016). It is likely that preovulatory aging disturbs various processes in the oocyte, all together resulting in a lower oocyte quality and impaired fertilization rates.

To analyze if early embryonic development is affected by preovulatory aging, we analyzed transcription during the first wave of zygotic genome activation (ZGA) at the 2-cell stage. This is a crucial step in preimplantation development required for successful maternal-to-embryo transition and insures development past the 2-cell stage (Minami *et al.* 2007). ZGA depends on maternal factors, which are known to be altered by preovulatory aging (Dankert *et al.* 2014). BrUTP incorporation was detected in the nucleoli of 2-cell embryos. In the nucleoli rRNA is synthesized, which is transcribed starting from the 2-cell stage (Knowland and Graham 1972). This is in line with a previous study that detected incorporation of BrUTP by RNA polymerase I, which transcribes rRNA in the nucleoli (Wansink *et al.* 1993).

Surprisingly, we found an increase in BrUTP incorporation in 2-cell embryos that derived from preovulatory-aged oocytes compared to controls, demonstrating a gain in transcription at this stage. This could point to a precocious activation of the embryonic genome prior to or early in the 2-cell stage, which has also been described for postovulatory aging (Xu *et al.* 1997). Postovulatory aging occurs when fertilization is delayed and can lead to transcriptional activation, resumption of the cell cycle, release of cortical granules and increased protein synthesis of factors required for ZGA in non-fertilized oocytes (Wang and Latham 1997; Xu *et al.* 1997). There is increasing evidence that many processes in the oocyte and during ZGA are regulated in a continuous, time-dependent manner. If events such as ovulation or fertilization are delayed, these processes do not adapt to this delay, which leads to a desynchronization and impaired oocyte quality. Wiekowski and colleagues (Wiekowski *et al.* 1991) described a “zygotic clock” that regulates ZGA independently of cell cycle events such as DNA replication, cell division or formation of the zygotic nucleus. More recently, indications for early recruitment of transcripts for protein translation during maturation of preovulatory-aged oocytes have supported this hypothesis (Dankert *et al.* 2014; Demond *et al.* 2016).

ZGA is a highly regulated process and depends on reprogramming of the parental genomes (Messerschmidt 2012). We previously found decreased transcript levels of maternal effect genes involved in epigenetic reprogramming of the embryonic genome (Dankert *et al.* 2014). Therefore, we investigated methylation levels of three imprinted genes (*Igf2r*, *Snrpn*, *H19*) and one unmethylated control gene (*Pou5f1*) at the 8-cell stage. Methylation levels of these genes should be maintained throughout epigenetic reprogramming in the preimplantation development. Indeed, we observed differential methylation for all investigated imprinted loci and hypomethylation of *Pou5f1*. We did not see an effect of preovulatory aging on methylation levels in these loci, indicating stable imprint maintenance. This is interesting considering the continuous debate whether assisted reproductive techniques (ART) may increase the risk for aberrant methylation and imprinting syndromes (Horsthemke and Ludwig 2005; Hiura *et al.* 2014; Lazaraviciute *et al.* 2014; Melamed *et al.* 2015). Due to the very low prevalence of imprinting syndromes, it is very difficult to determine whether the increased prevalence of these syndromes in children conceived with ART is actually caused by the procedure or whether there is a genetic base in the patient's cohort (Lazaraviciute *et al.* 2014). In mice, a direct effect of in vitro fertilization on methylation of imprinted genes has been described and there is evidence that superovulation might be the underlying cause (Fauque *et al.* 2007; Sato *et al.* 2007; Market-Velker *et al.* 2010; de Waal *et al.* 2012; Fortier *et al.* 2014; De Waal *et al.* 2015; Huffman *et al.* 2015). We did not see such effects, which might be due to a dose-dependent effect of hormone treatment (Market-Velker *et al.* 2010). Our present study is not the only one though that describes stable imprint maintenance after hormonal treatment. Denomme et al. (Denomme *et al.* 2011) observed no effect on DNA methylation of imprinted genes in mouse oocytes after superovulation. It is likely that there is an cumulative effect of not only superovulation but also other factors such as culture media that may cause the increased risk for imprint defects during ART (De Waal *et al.* 2015).

In line with previous findings, we observed that different developmental processes are affected in different ways during preovulatory aging. The present results show stable DNA methylation in embryos after preovulatory aging at all three analyzed loci. However, impaired fertilization rates and altered embryonic genome activation indicate that preovulatory aging affects maternal factors, which may lead to a reduced developmental competence. Embryonic genome activation seems to be not (exclusively) timed by ovulation but at least partly regulated by events in the growing or maturing oocyte. A delay in ovulation desynchronizes physiological processes in the oocyte, which may consequently affect the embryonic program resulting in potentially long-lasting effects for the offspring. In a broader sense, this might be important for reproductive health in humans, since cycle irregularities are associated with a number of conditions such as polycystic ovary syndrome (PCOS) or primary ovary insufficiency.

## Material and methods

### Ethics statement

This study was carried out in strict accordance with the recommendations in the Guide for the Care and Use of Laboratory Animals of the German Government. The protocol was approved by the Committee on the Ethics of Animal Experiments of the responsible authorities (Landesamt für Natur, Umwelt und Verbraucherschutz, LANUV AZ 84-02.04.2011.A374). All animals were kept under standard conditions (food and water *ad libitum*, 12:12 h light and dark cycles) in the Central Animal Laboratory of the University Hospital Essen. Animals were acquired from the breeding colonies of the animal facility.

### Mating and embryo collection after preovulatory oocyte aging

Preovulatory oocyte aging was induced by delaying ovulation, leading to prolonged oocyte maturation and therefore causing oocyte overripeness (Bittner *et al.* 2011). Follicle growth was stimulated with pregnant mare serum gonadotropin (PMSG) and superovulation was induced with human chorionic gonadotropin (hCG) 48 hours later in control animals. From the start of hormonal treatment, the GnRH antagonist cetrorelix (Cetrotide, Merck-Serono) was applied every day, in order to delay ovulation for 3 days before hCG injection and inducing preovulatory aging as described previously (Dankert *et al.* 2014).

To avoid inbred depression and to increase embryo yield for BrUTP incorporation analysis, C57Bl/6J females were mated with C57Bl/6J x CBA/Ca hybrid males after hCG injection. Females with vaginal plug on the next morning were sacrificed by cervical dislocation 24 hours post hCG stimulation. Two-cell embryos were collected in M2-Medium (Sigma-Aldrich) by flushing of the oviduct and immediately used for investigation of BrUTP incorporation.

For DNA methylation bisulfite-sequencing analysis, C57Bl/6J females were mated with CAST/EiJ males. These two mouse strains contain single nucleotide polymorphisms (SNPs) in the analyzed loci, allowing discrimination between the maternal and paternal allele during sequencing. Two-cell embryos were collected as described above and further cultured in KSOM medium (GlobalStem) under mineral oil for 24 hours at 37 °C and 5% CO2 until the 8-cell stage. Each single 8-cell embryo was collected in 10 μL PBS and stored at -20 °C until further use.

### BrUTP incorporation and immunofluorescence analysis

Embryonic genome activation was analyzed by measuring the incorporation of 5-bromouridine-5′-triphosphate (BrUTP) in *de novo* synthesized RNA. The BrUTP incorporation assay was performed as described previously (Aoki *et al.* 1997). Briefly, 2-cell embryos were incubated in BrUTP for 10 minutes. Incorporation of BrUTP was visualized using a FITC-labeled mouse anti-BrdU monoclonal antibody (1:50, Roche Diagnostic). To validate the BrUTP antibody specificity, transcriptionally quiescent oocytes were used as negative controls and mouse anti-human CEACAM1 antibody as an isotype control. The 2-cell embryos were mounted in Vectashield solution with 4′,6-diamidino-2-phenylindole (DAPI; Vector Laboratories). The fluorescent antibody staining was detected with a Leica DM4000B microscope. The fluorescent signal intensity was quantified with Image J software 1.48v.

### Preparation of bisulfite sequencing libraries

DNA from single 8-cell embryos was extracted and treated with bisulfite using the EZ DNA Methylation-Direct Kit (Zymo Research) according the manufacturer’s protocol and eluted in 10 μl M-Elution Buffer. To generate locus-specific amplicon libraries a nested PCR approach was used. PCR and sequencing primers were described previously by El Hajj et al. (2011; Table S1). Three imprinted genes (*H19*, *Snrpn* and *Igf2r*) were analyzed and *Pou5f1* was used as a control gene that should be expressed and have no methylation at the 8-cell stage. The analyzed amplicons contained between 4 and 14 CpGs and all amplified loci contained strain-specific SNPs to distinguish between the maternal and paternal allele (Table S1). This is important, since bisulfite treatment destroys DNA and with only 8 cells as starting material there is a high risk of a bias towards either the maternal or paternal allele. Bisulfite converted DNA was first amplified in a multiplex PCR, where outer primers for all 4 loci were mixed, using the Multiplex PCR Kit (Qiagen). For each embryo, a 25 μl reaction was set up, containing 12.5 μl 2× Multiplex PCR Mastermix, 2.5 μl 10× Primer Mix (2 μM per primer) and 10 μL bisulfite treated DNA template. After initial denaturation for 15 min at 95 °C the DNA was amplified in 30 cycles of 30 s at 94 °C, 90 s at 52 °C and 45 s at 72 °C. Final extension occurred at 72 °C for 10 min. This was followed by a second, inner PCR step with tagged gene specific primers. For each gene 12.5 μl 2x HotStarTaq Mastermix (Qiagen) was mixed with gene specific 2.5 μL FWD and 2.5 μl REV primers (5 μM each) and 3.0 μl outer PCR product. 4.5 μl water were added to reach a final volume of 25 μl. After initial denaturation for 15 min at 95 °C, a touchdown PCR protocol was carried out to increase primer specificity. Starting at an annealing temperature of 64 °C, the annealing temperature was reduced in 14 cycles to 57 °C by decreasing the temperature 0.5 °C in each cycle. This was followed by additional 25 cycles of 45 s at 72 °C, 30 s at 57 °C, 45 s at 72 °C and final extension for 10 min at 72 °C. Samples were sequenced on the Roche/454 GS junior system. Before sequencing, a third PCR was used to add a sample-specific barcode. This consisted of a sample-specific multiplex identifier (MID) and the universal linker tag that contains a sequence complementary to the tagged primers from the second PCR and furthermore includes 454 adaptor sequences and a key. A 50 μl PCR reaction was set up with 25 μl 2x HotStarTaq Mastermix, 0.5 μl FWD and 0.5 μl REV primer (20 μM each), 3 μl PCR product from the second PCR and filled up with water. PCR conditions were as follows: 10 min denaturation at 95 °C, then 35 cycles with 20 s at 95 °C and 30 s at 72 °C and a final elongation step for 7 min at 72 °C.

### Deep bisulfite sequencing using the Roche/454 Genome Sequencer

Sample preparation and sequencing on the Roche/454 GS junior were carried out as previously described (Beygo *et al.* 2013). In short, amplicons from the third PCR were purified using the Agencourt AMPure XP Beads system (Beckman Coulter) following the recommendations by Roche. Samples were quantified using the NanoDrop ND-1000 Spectrophotometer (ThermoScientific). Libraries were diluted, pooled and clonally amplified in an emulsion PCR (emPCR). Sequencing on the Roche/454 GS junior sequencer was carried out according to manufacturer’s instructions (Roche emPCR Amplification Method Manual – Lib-A and Roche Sequencing Method Manual).

Data analysis was conducted using the Amplikyzer Software (Rahmann *et al.* 2013) with default settings based on the ⋅ssf files generated by the 454/Roche GS Junior system. The methylation level of the maternal and paternal allele was determined using SNPs. Due to the destruction of DNA during bisulfite treatment and the low input amount it was not always possible to obtain reads for both the maternal and the paternal allele. In the case that both alleles were present for analysis the mean methylation level of the two alleles was calculated, which was expected to be 50% for imprinted genes and 0% for the *Pou5f1* control gene.

### Statistical analysis

Statistical analysis was performed using SigmaPlot version 12.5 (Systat Software, Inc). Normal distribution of the data was analyzed with the Shapiro-Wilk-test. Normal distributed data was compared with a two-sided student's *t*-test. When the Shapiro-Wilk-test failed a Mann-Whitney U-test was conducted to calculate the difference between two data sets. Fertilization rates were analyzed as number of embryos per mouse, with mouse numbers determining sample size. For analysis of BrUTP incorporation and DNA methylation embryo numbers used for the study were taken as sample size. A post-hoc power analysis of the DNA methylation data was conducted using G*Power version 3.1.9.2. The α-level was set at 0.05 to determine statistically significant differences. All data are shown as mean ± SEM.

## Acknowledgement

We thank Laura Steenpass, Jasmin Beygo and Lars Maßhöfer for their help during the establishment of the bisulfite-sequencing protocol and Sabine Kaya for performing the sequencing analysis. We are grateful to Ulrich Zechner of the University of Mainz, Germany, who kindly provided the breeding founder animals for the CAST/EiJ strain. This work was funded by DFG Research Grants: HO 949/21-1 and GR 1138/12-1.

## Supporting information

**S1 Figure: Comparative analysis of DNA methylation levels at the *Igf2r* locus of control embryos.** Indicated are the mean DNA methylation levels of the entire locus per embryo of the maternal allel **(A)** and the paternal allel **(B)**. The white numbers indicate methylation levels of single CpGs. The colors represent a scale from red (100% methylation) to blue (0% methylation).

**S2 Figure: Comparative analysis of DNA methylation levels at the** ***Igf2r*** **locus of preovulatory-aged (PreOA) embryos.**

**S3 Figure: Comparative analysis of DNA methylation levels at the** ***Snrpn*** **locus of control embryos.**

**S4 Figure: Comparative analysis of DNA methylation levels at the** ***Snrpn*** **locus of preovulatory-aged (PreOA) embryos.**

**S5 Figure: Comparative analysis of DNA methylation levels at the** ***H19*** **locus of control embryos.**

**S6 Figure: Comparative analysis of DNA methylation levels at the** ***H19*** **locus of preovulatory-aged (PreOA) embryos.**

**S7 Figure: Comparative analysis of DNA methylation levels at the** ***Pou5f1*** **locus of control embryos.**

**S8 Figure: Comparative analysis of DNA methylation levels at the** ***Pou5f1*** **locus of preovulatory-aged (PreOA) embryos.**

**S1 Table: Primers for DNA methylation analysis of embryos.**

